# Saccadic modulation of neural excitability in auditory areas of the neocortex

**DOI:** 10.1101/2022.05.24.493336

**Authors:** Marcin Leszczynski, Stephan Bickel, Maximilian Nentwich, Brian E. Russ, Lucas Parra, Peter Lakatos, Ashesh Mehta, Charles E. Schroeder

**Author notes:** Correspondence: Dr. Marcin Leszczynski and Dr. C.E. Schroeder, Department of Psychiatry, Columbia University College of Physicians and Surgeons, 1051 Riverside Drive Kolb Annex Rm 561, New York, NY 10032.

## Abstract

**Summary:** In natural “active” vision, humans and other primates use eye movements (saccades) to sample bits of information from visual scenes. In this process, nonretinal signals linked to saccades shift visual cortical neurons to a high excitability state as each saccade ends. The extent of this saccadic modulation outside of the visual system is unknown. Here, we show that during natural viewing, saccades modulate excitability in numerous auditory cortical areas, with a pattern complementary to that seen in visual areas. Bi-directional functional connectivity patterns suggest that these effects may arise from regions involved in saccade generation. By using saccadic signals to yoke excitability states in auditory areas to those in visual areas, the brain can improve information processing in complex natural settings.

## Introduction

Humans and other primates actively sample the visual environment by systematically shifting eye gaze several times per second^1^. These rapid “saccadic” eye movements trigger volleys of retinal input that drive neural activity along the visual pathway^2–4^ from the lateral geniculate nucleus^5,6^ through several stages of the visual cortex^7–9^ to the medial temporal lobe and the hippocampus^10–12^. The effects of saccades in the dark^7,13^, and those noted in studies that minimize retinal inputs initiated by saccades^14,15^, support the idea that the gain of retinal input processing in visual areas is modulated by non-retinal signals, generated in parallel to saccades^16^. Saccadic perturbation of neural excitability by both driving and modulatory inputs occurs rhythmically at the 3-5 Hz saccadic rate – a key physiological signature of active sensing^17–19^. Saccade-related reset of neuronal oscillations in this “theta” range provides a possible mechanism for modulation of neuronal activity^17,18^. The Frontal Eye Fields (FEF) are involved in preparing and generating saccadic eye movements^20^. The FEF also coordinate top-down attention allocation and modulate neural activity in the visual system through network connectivity^21^ oscillating in gamma and beta ranges^22^.

Intriguingly, several studies suggest that the influence of saccades on neural activity extends beyond the visual system. Sensitivity to static eye position has been noted in both subcortical and cortical auditory pathway structures^23,24^ and dynamic saccadic modulation has been observed in the non-visual nuclei of the anterior thalamus^25^ as well as in the auditory cortex^26^. Notably, in both monkeys and humans, the eardrum moves in synchrony with saccades^27^. However, the extent of saccadic modulation across auditory cortical regions has not been determined, and the brain circuits projecting these effects into auditory areas have not been identified. Here, we combined eye tracking during natural free viewing with simultaneous intracranial recordings from widespread brain regions in surgical epilepsy patients to address these questions.

We have several key findings. First, saccadic modulation of neural excitability is much more widespread than previously recognized. In the auditory selective networks (ASNs; those responding preferentially to auditory stimuli), saccades modulate low frequency phase and power with distinct anatomical and spectrotemporal profiles. Phase modulations were observed primarily in higher order areas, while power modulations occurred in lower level auditory areas. Second, we found that Broadband High-frequency Activity, or BHA^28^ was increased towards the end of the fixation period and during the ensuing saccade, suggesting the local neuronal excitability in the ASNs is transiently enhanced with a time-course complementary to that of the visual system. Finally, we observed that theta/alpha and beta range patterns of functional connectivity between FEF and the ASNs changed magnitude and direction over the saccade-fixation cycle. The connectivity profile suggests that excitability in the ASNs is modulated by top-down projections from the FEF. These results support a model in which saccadic modulation of neuronal excitability over a wide domain of auditory neocortical areas may complement similar modulation of excitability the visual pathways to stabilize information processing during rhythmic saccadic sampling in complex natural environments.

## Results

### Saccades modulate neural excitability in the auditory selective networks (ASNs)

We investigated the influence of saccades (Fig. 1A) on neural activity in ASNs by analyzing the dynamics of field potentials across the saccade-fixation cycle. Using standard auditory and visual localizers in 9 surgical epilepsy patients, we identified 220 auditory-selective sites (i.e., channels that responded to sounds but gave no detectable response to visual stimulation; Fig 1B). Next, patients performed a free-viewing task in which they explored a set of 80 images each presented for the duration of 6 sec in the absence of any auditory stimulation. Using simultaneous eye tracking, we defined time points corresponding to fixation onset and locked the analyses of electrophysiological signals relative to these time points (Fig. 1C). As in our earlier studies^7,15,25,28^, we focus on neural events related to the end of the saccade (fixation onset) as this is an event with clear perceptual relevance; i.e., the point at which retinal inflow drives the ascending visual pathways. On average, participants made 1054 fixations (standard deviation of 245) with the median duration of fixation of 239 ms (Inter-quartile-range of 52 ms). The resulting 4 Hz rate of saccadic exploration during natural free viewing is typical for both human and non-human primates^10,11,15,25^.

**Figure 1.**
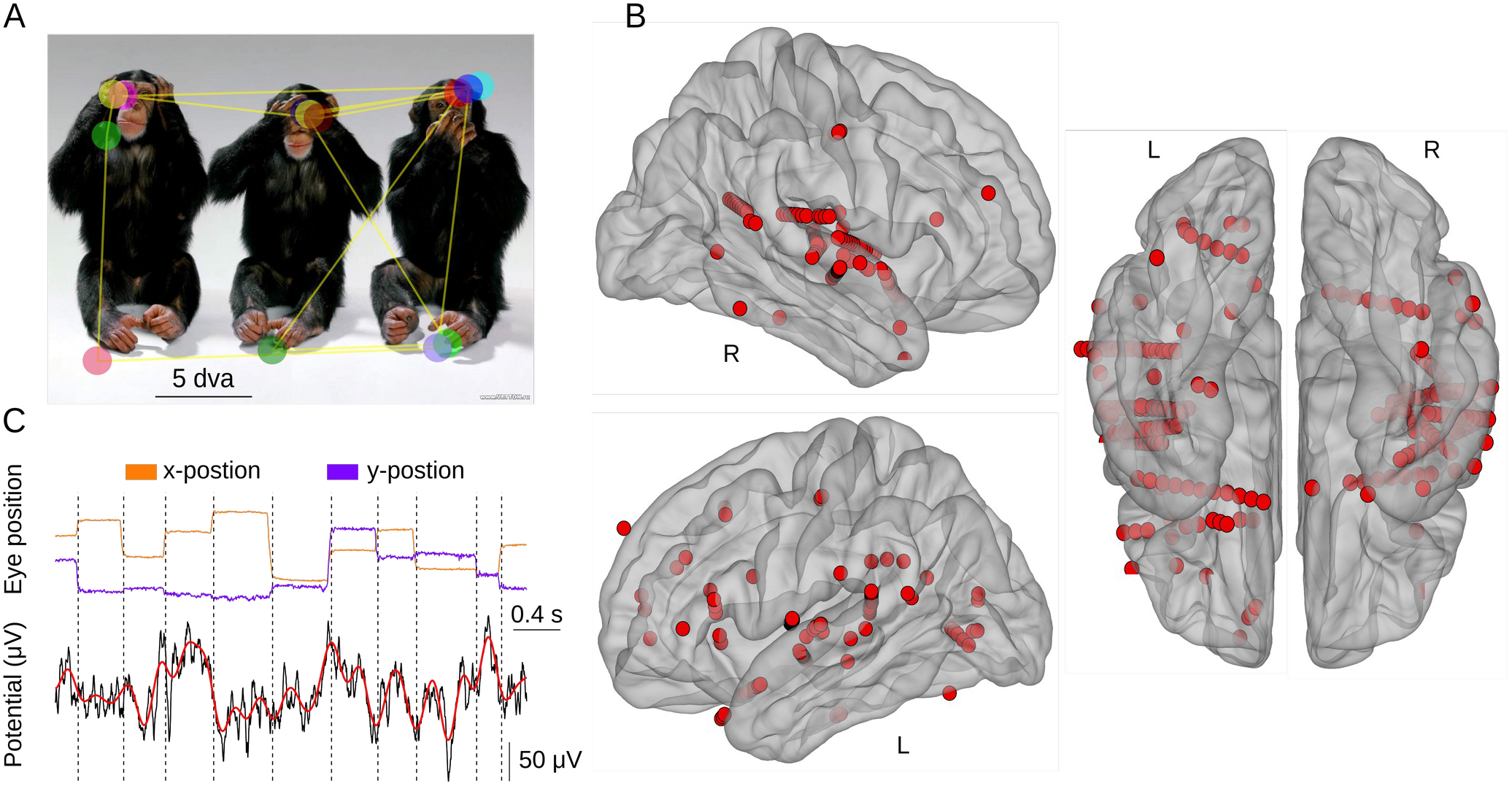
Task, example of the field potentials and the anatomical distribution of auditory selective channels. **(A)** Example image with superimposed trace of eye movements (yellow lines) and fixations (colored circles) during free viewing. **(B)** Anatomical distribution of auditory selective electrodes (N = 220 electrodes across 9 patients), that in the localizer condition respond with an increased BHA to auditory stimuli and no detectable response to visual stimuli. Electrodes from all patients were transformed onto an average brain. Left panels: lateral view. Right panels: inferior view (R – right; L – left). **(C)** Example eye tracking traces (x- and y-position of the eye presented in orange and violet, respectively) and a corresponding intracranial field potential from one electrode. Vertical lines mark points of fixation onset. The lower panel presents raw field potential (black) and low-passed filtered at 5 Hz signal (the maximum saccade rate; red).

To investigate saccadic modulation of neural activity in the ASNs, we studied fixation-related changes in neural dynamics, as indexed by inter-trial phase coherence (ITC) and power in low frequencies (<30 Hz). We also studied fixation-related changes in Broadband High frequency Activity (BHA), which reflect neuronal processes, that are separable from, but highly correlated with neuronal firing^28^.

We found two spectrotemporal ITC components: 1) an early broad-band component centered at 80-100 ms after fixation onset in the beta frequency range, extending down into alpha and theta ranges (Fig 2A); 2) a later band-limited component that was strongest around 200 ms after fixation onset and limited to the theta range, which corresponds to the rate of eye movement exploration (i.e., 3-5 Hz; Fig 2A). Quantifying and directly comparing the magnitude of ITC in early (0-100 ms) and late (100-400 ms) post-fixation time windows relative to the pre-stimulus interval (−400 to 0 ms) confirmed the above impressions (see Fig. 2A; Wilcoxon tests; p < 0.05; unless stated different all results are controlled for multiple comparisons with Benjamini & Hochberg/Yekutieli false discovery rate procedure; see Methods). To ensure that these results are not biased by weak visual evoked responses in a subset of channels, we directly quantified fixation-locked evoked responses and differentiated a subgroup of channels with strong audio and weak visual evoked responses (i.e., below the initial detection threshold). The control analyses confirm that the effects are unchanged when weak fixation-locked evoked response sites are excluded (see Supplemental Information).

**Figure 2.**
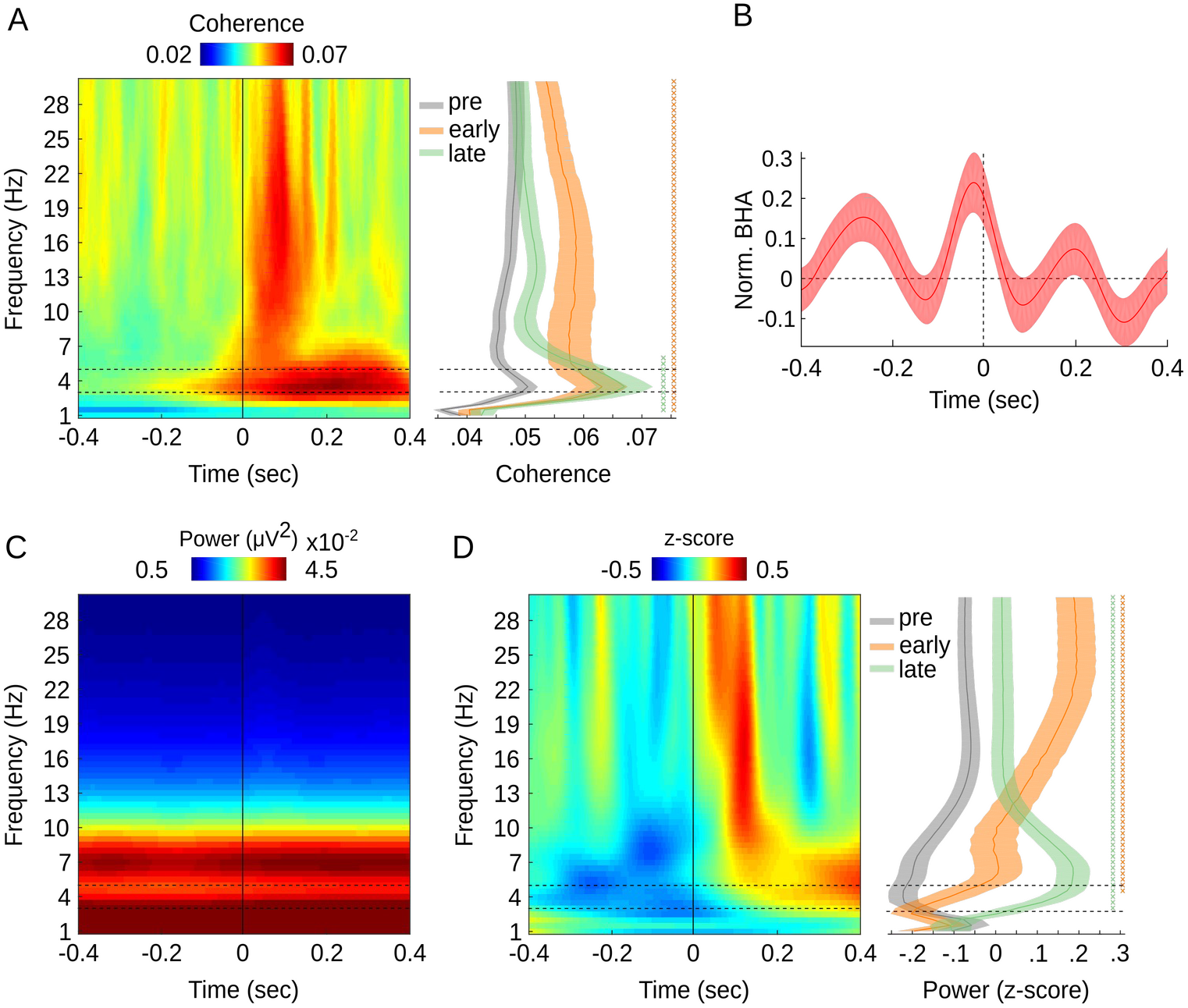
Fixation-locked neural activity in the ASNs. **(A)** Color map shows average fixation-locked ITC (N = 220 channels, 9 patients; x-axis: time, y-axis: frequency). Vertical line indicates fixation onset. Horizontal lines bracket the frequency range of saccades (3 – 5 Hz). Right panel shows frequency distributions of ITC averaged within three time windows relative to fixation-onset: pre-fixation (gray; -400-0 ms), early-post (0-100 ms; orange) and late-post (100-400 ms; green). Shading reflects standard error of the mean (SEM). **(B)** Z-scored BHA locked to fixation onset aggregated across all ASNs. **(C, D)** Color maps show fixation-locked power without normalization (C) and normalized (D) by z-scoring each frequency separately to remove the 1/f component (x-axis: time; y-axis: frequency). The right panel (in D) shows frequency range of power fluctuations within three time windows relative to fixation-onset (as in panel A). All results are controlled for multiple comparisons with Benjamini & Hochberg/Yekutieli false discovery rate procedure. “X” symbols in right panels A, D indicate p-values < 0.05 separately for the comparison between pre-vs. early- and late-post fixation time window (orange and green, respectively).

To test the hypothesis that saccades modulate neural excitability in the ASNs, we measured the magnitude of the BHA signal from the pre-to the post-fixation time intervals. We found that overall post-fixation (0-75 ms) BHA magnitude was reduced (z = 3.1, p = 0.001; N = 220; Fig. 2B) compared to the pre-fixation interval (−75-0 ms). To rule out the possibility that the pre-fixation BHA increases spuriously originate from extraocular muscle activity^29^ we analyzed saccade-related field potentials. We reproduced our results after eliminating any channels that showed “spike” morphology similar to that of extraocular muscles (see Fig. S1 and Supplemental Information). In sum, BHA power fluctuation in the ASNs is synchronized with the rhythm of visual exploration and increased in the pre-as compared to the post-fixation time interval.

Finally, we examined spectral power modulations in low frequencies (<30 Hz; Fig. 2C-D). Because we used electrode sites that respond selectively to the auditory classifier, it is not surprising that we did not observe an effect typical for visual system, i.e., a fixation-locked evoked response. Instead we found a prominent narrow-band power increase in the range just above the rate of visual exploration (7-10 Hz). Spectral power decreased in the pre-fixation period compared to both early and late post-fixation time windows. While we observed weak power modulations in a broad range of frequencies, power modulations in both time windows were centered at about 7 Hz (Fig. 2C-D). In sum, our findings show that neural activity in the ASNs is synchronized with the rhythm of saccades. Specifically, we find 1) increased ITC at the rate of visual exploration; 2) alpha-band power decrease and 3) that BHA fluctuation is synchronized to saccades with a magnitude increase near the end of fixation and during the ensuing saccade. All of these findings suggest that saccades modulate neural excitability in the ASNs, such that it is highest near the end of fixation period and during the ensuing saccade.

### Auditory responses are transiently enhanced during saccades

Next, we investigated modulation of auditory evoked responses in the ASNs across the saccade-fixation cycle. We reasoned that momentarily increased neural excitability at any point in the saccade-fixation cycle would increase the strength of concomitant neural responses to auditory stimulation. To test this idea, we recorded patients while they viewed movies with and without a continuous audio stream (Methods), as well as the static scenes; this was done in 8 of the 9 patients reported above (N = 187 ASNs channels). We normalized fixation-locked BHA time courses by z-scoring each individual condition so that they were all centered to a common value. This enabled us to directly compare time courses of fixation-locked BHA across the three conditions: free-viewing of movies 1) with, 2) without audio and 3) static images.

The overall dynamics of fixation-locked BHA during free viewing of movies (both with and without the audio stream) were similar to those observed during free viewing of static images – BHA peaked prior to fixation onset followed by a dip around 100 ms after fixation onset (Fig. 3A). Importantly, we found that BHA in the time interval prior to fixation onset was elevated during viewing of movies with audio (Fig. 3A,B) as compared to both movies with no audio (z = 3.9; p < 0.001; N = 187) and static images (z = 2.4; p = 0.01; N = 187). There was no detectable difference in BHA magnitude between conditions with no auditory stimulation (z = 1.81, p = 0.07; N = 187) or between any of the conditions in the time window following fixation onset (all z < 1.6; all p > 0.1; N = 187). We reproduced these effects in a control analysis in which we excluded weak visual response channels, as discussed above (Fig. S6A,B). To ensure these effects are not due simply to the presence or absence of the auditory input we confirmed that the intensity of the auditory stimulus stream did not differ between pre-(−0.1-0 sec) and post-fixation (0-0.1 sec) time windows; this held true whether we considered raw audio input or the envelope of the wide band filtered auditory stream (both z < 1.26, p > 0.20; N = 8; patients; Wilcoxon test). Similarly, there was no detectable difference in wider time windows of +/-0.4 and +/-0.6 sec (all z < 1.12; all p > 0.26; N = 8; patients; Wilcoxon test). Altogether, these results show that during natural free viewing, neural response to the auditory stimulation in the ASNs is transiently amplified just prior to a given fixation onset (i.e., late in the fixation period and during the ensuing saccade). Because auditory stimulation was comparable throughout the saccade-fixation cycle, we conclude that the BHA enhancement results from increased excitability during this part of the fixation-saccade cycle.

**Figure 3.**
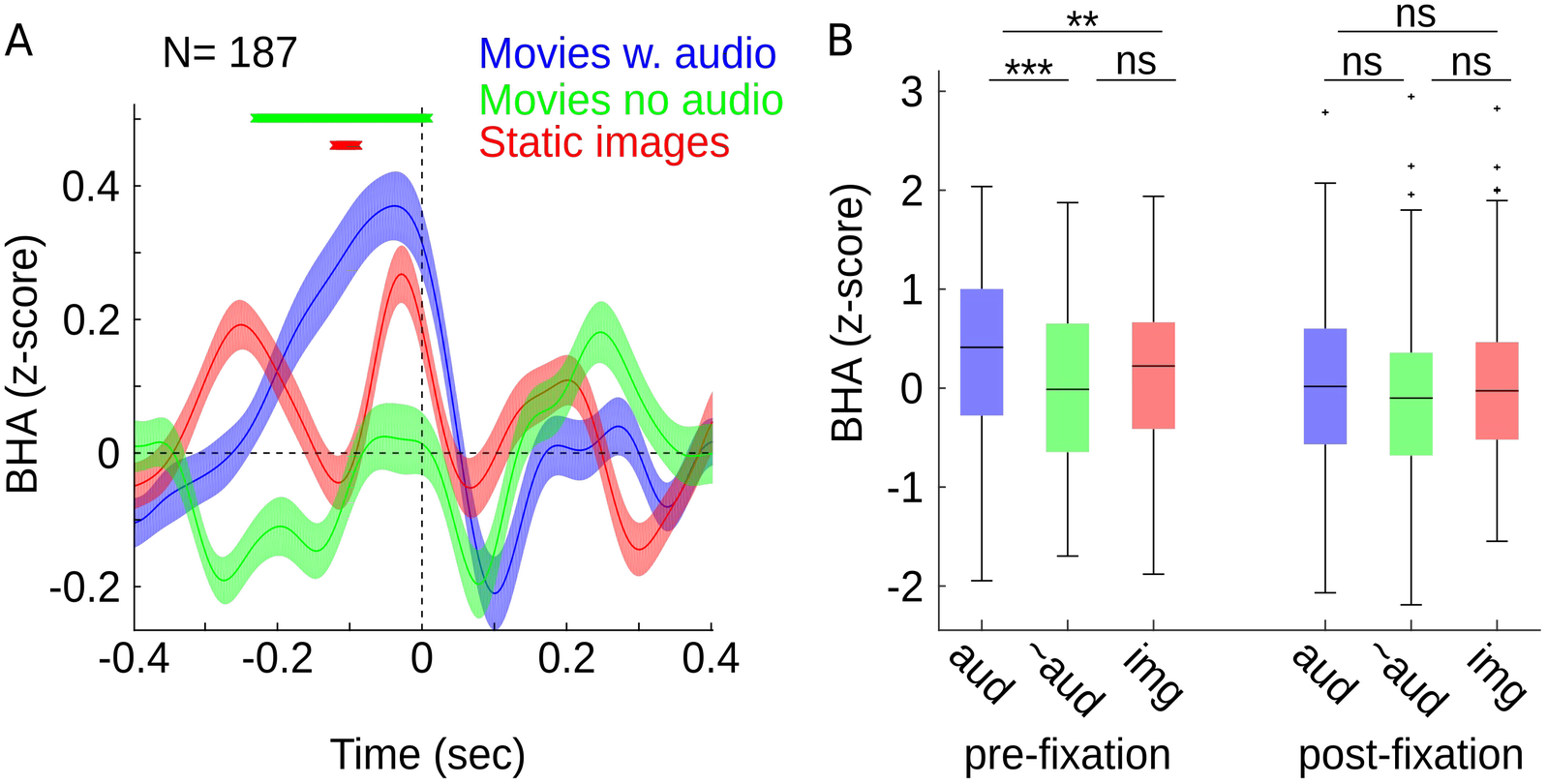
Saccades transiently amplify neural responses to auditory stimulation. **(A)** Fixation-locked BHA (z-scored) in three viewing conditions: free viewing of movies with audio stream (blue), movies with no audio (green) and free viewing of static images (red). Data are aggregated across all 187 ASNs. This number is lower than reported in analyses above because only 8 out 9 patients performed all three viewing conditions. X-axis: time relative to fixation onset; Y-axis: magnitude of BHA modulation. Shading reflects SEM. “X” symbols in panel A indicate p-values < 0.05 (controlled for multiple comparisons) separately for two a priori comparisons: movies with audio vs. movies with no audio (green) and movies with audio vs. static images (red). **(B)** Box plots show the magnitude of fixation-locked BHA in pre-(−100:0 ms) and post-fixation time windows (0:100 ms). Box plots indicate 25^th^, median and 75^th^ percentile, whiskers extend to extreme values not considered outliers while outliers are marked with crosses. Star symbols indicate p-values: *,**,*** signifies p < 0.05, 0.01, 0.005, respectively; ns signifies p > 0.05.

### Saccades organize bidirectional communication between the FEF and ASNs

The finding that saccades modulate neural activity in the ASNs suggests that like visual networks, ASNs are modulated by a top-down influence from regions involved in saccade generation, such as the FEF. To test this possibility, we examined the direction of network interactions between the FEF and ASNs using Phase Slope Index (PSI)^30^. Specifically, we focused on channels in the Superior Temporal Gyrus (STG, 54 channels across 4 patients with simultaneous FEF recordings) because this region of interest was present in all patients with FEF electrodes and previous anatomy studies found that neurons in the FEF (particularly the rostral part of the FEF) project primarily to the STG^31^. It is however worth noting that the results are similar whether we consider all auditory selective channels or only STG sites (Fig. S8).

If FEF increases neural excitability in the auditory system prior to fixation onset, we expect to see primarily top-down directional network interactions from the FEF to STG towards the end of fixation. This top-down interaction would be reversed to a primarily bottom-up direction after fixation onset when sensory signals propagate up sensory hierarchies to regions programming the next eye movement. Because the sign of the PSI reflects the direction of network interactions, following previous studies^30^, we tested the magnitude of PSI against a null hypothesis that PSI is not different from zero (see Methods).

We found significant directional network interactions in theta/alpha (3-12 Hz) and two beta frequencies (lower-12-20 Hz and higher-beta 22-27 Hz; Fig. 4C-D). Importantly, we observed that these directional network interactions were dynamically changing in time across the saccade-fixation cycle. Bottom-up interaction from the STG to FEF was in theta/alpha- and higher beta frequencies while top-down from the FEF to STG in lower beta frequency (Fig. 4C). We also found that the theta/alpha bottom-up interaction was significant throughout the entire saccade-fixation cycle with a noticeable frequency expansion and dip of the effect at fixation onset. Both lower and higher beta range effects depended on the temporal position within the saccade-fixation cycle. Importantly, the lower beta top-down influence was strongest late in fixation and during saccades. The higher beta bottom-up influence was in turn strongest right after fixation onset (Fig. 4C-D). Together, these results suggest that top-down and bottom-up interactions between the FEF and the STG are multiplexed in frequency and dynamically change across the saccade-fixation with a burst of top-down interactions concentrated in lower beta frequency range late in fixation period and during the ensuing saccade.

**Figure 4.**
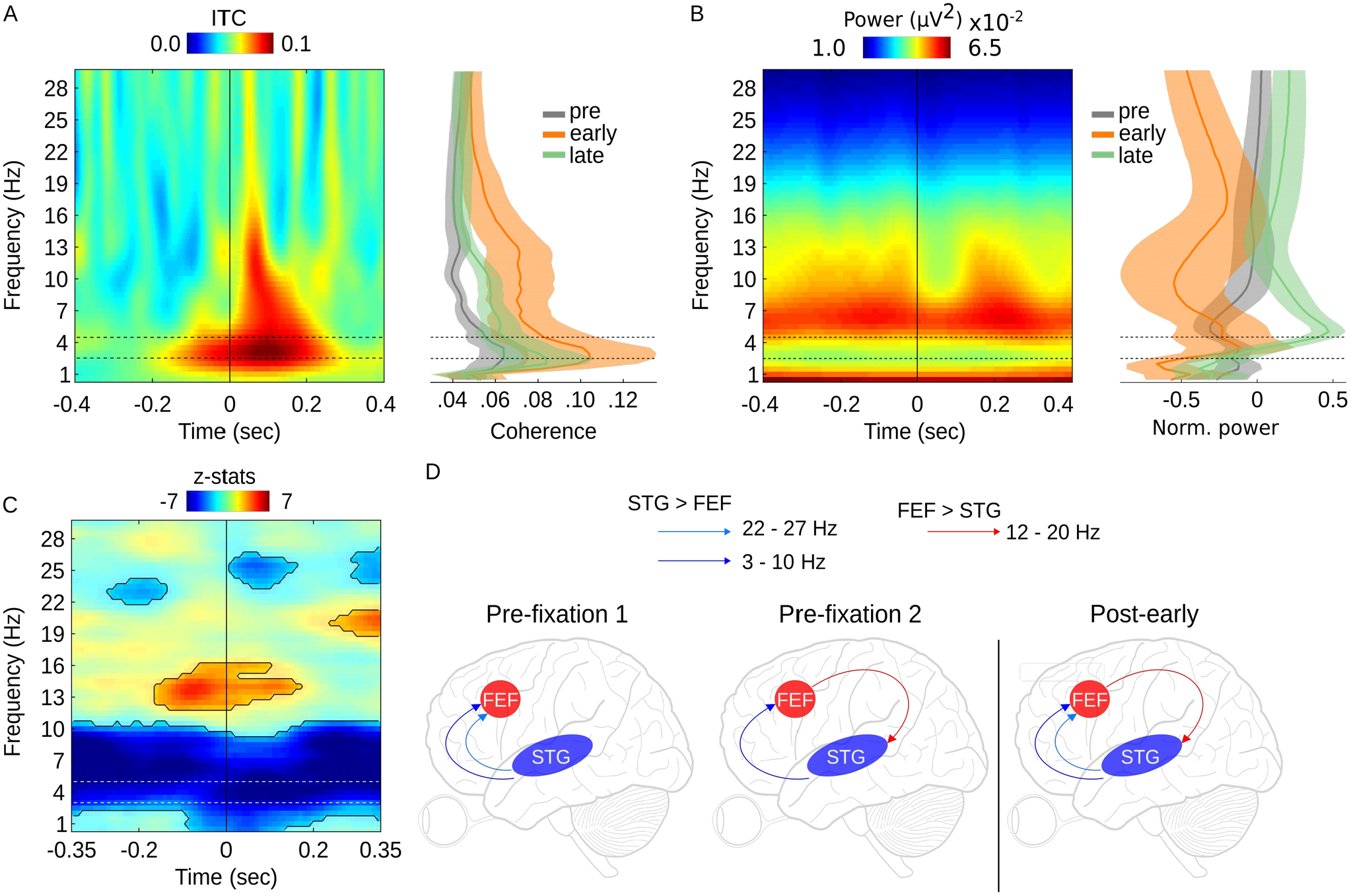
Fixation-locked network connectivity between FEF and STG. Color maps show averaged fixation-locked ITC (**A**; N = 8 channels, 4 patients; x-axis: time, y-axis: frequency) and power **(B)** in FEF. Right panels show frequency distributions of ITC (A), power (B) in three time windows relative to fixation-onset: pre-fixation (−400:0 ms; gray), early-post (0:100 ms; orange), late-post (100:400 ms; green). Shading reflects SEM. **(C)** Fixation-locked z-statistics from a Wilcoxon test comparing PSI against a null hypothesis (i.e., PSI not different from zero). Negative values (blue) indicate points when STG leads FEF (i.e., bottom-up), positive values (red) indicate reversed (i.e., top-down) direction. Contours depict significant time-frequency points (p < 0.05 controlled for multiple comparisons). **(D)** Schematic representation of the network interactions as observed in C. Theta/alpha-(dark blue) and upper beta-band (light blue) interaction from the STG to FEF at about 200 ms prior to fixation (pre-fixation 1). The upper beta activity was attenuated during the saccade and replaced by top-down interaction in lower beta (pre-fixation 2; red), which occurred with theta/alpha-band bottom-up activity (dark blue). After fixation onset, all three activities lasted for about 200 ms (post-early). Vertical line marks fixation onset. Arrows indicate direction of network interaction.

## Discussion

Combining eye tracking and iEEG in surgical epilepsy patients during naturalistic free viewing, we found that the saccade-fixation cycle is associated with systematic modulation of neural excitability across multiple classical auditory and speech processing regions. We observed fixation-locked phase concentration at the rate of saccades as well as sustained power concentration in the lower part of the alpha spectrum (∼7-10 Hz) that dropped before fixation onset. These effects showed distinctive anatomical distributions with phase effects linked to higher order speech regions and power effects observed in primary and secondary auditory regions. Broadband High-frequency Activity (BHA), a proxy for local neuronal excitability, was increased late in the fixation period and during the ensuing saccade, and neural responses to auditory stimulation were transiently enhanced during this time frame. Finally, we observed dynamic bi-directional network interactions between the FEF and STG in theta/alpha, higher beta and lower beta frequencies with strongest top-down interaction measured in the lower beta frequency range late in fixation period and during saccades. This overall pattern of results indicates that during natural active vision, local neural activity in auditory system is dynamically modulated over the course of the saccade-fixation cycle, likely through long-range interactions with networks involved in generation of saccades.

### Saccadic modulation beyond the visual system

Previous studies found that neural ensembles in the visual system are transiently reorganized to a high-excitability state after fixation onset (i.e., during the time when visual inputs propagate in the visual cortex) which enhances neural responses to sensory inputs^7–9^. Similarly, in rodents, natural active sensing boosts neural activity in primary sensory areas^32–34^. Here, we show that a critical form of active sensing in humans, saccadic sampling of the visual scene, modulates neural activity across a much larger network than previously recognized. This network extends into the classical auditory and speech processing regions with distinct effects in the lower and higher order areas.

### Saccadic modulation in relation to auditory processes

We noted a contrast between the increased phase coherence starting around 80 ms after fixation onset in higher order auditory areas in the frontal and parietal regions, and the BHA and alpha power modulations that are strongest before fixation onset in lower level auditory areas. This contrast raises possibility that there are two mechanisms operating in synchrony with saccades in the auditory system. On the one hand, phase coherence effects may reflect retinal inputs propagating from the visual system to higher level auditory system, related to multisensory integration (see e.g.,^35^). On the other hand, excitability, as indexed here by BHA magnitude and alpha power modulations, appears highest *before* fixation onset (see also^26^) and likely reflects a saccade related preparatory influence. It is possible that the elevation of excitability across the auditory system before fixation onset reflects a momentary deployment of auditory attention. Psychophysical findings suggest that as in vision^36^, auditory attention is spatially and temporally linked to the saccade-fixation cycle^37–39^. The time course of saccadic influences on auditory discrimination shows that saccade’s impact starts prior to movement onset and peaks while the eyes are moving^38^, which corresponds with the timing of BHA and alpha power modulation we observe as well as that of multiunit activity modulation in monkey A1^26^. Our results, together with those of prior psychophysical studies, suggest that increased pre-fixation excitability modulation may have a strong attentional component. In support of this idea, we observed increased directional network interactions from the FEF to the auditory system late in fixation period and during the ensuing saccade. It is possible that this top-down signal from the FEF modulates excitability in the auditory system implementing a physiological mechanism for transiently enhancing auditory attention in a way that complements saccade-related enhancement of visual attention^21,22,40^. If auditory attention is yoked to saccades, one would predict that it samples auditory space periodically at the rate of saccades. Indeed, similar to vision^41,42^ auditory attention has been recently suggested to operate in a periodic mode^43,44^.

Whether or not the signal modulating neural activity in auditory areas during saccades indeed reflects attention, it clearly does influence auditory processing and likely plays a role in natural active sensing. What might this role be? One possibility is that audition, and possibly other senses, compensate for reduced visual sensitivity during saccadic suppression. This idea builds on the observation that momentary increases in excitability and enhancement of auditory attention coincide with reduced neural response magnitude and depression of sensitivity in the visual system^36^. Such a mechanism may be particularly useful to compensate for loss of sensitivity in the magnocellular visual pathway which is strongly suppressed during saccades^36^. Given the preferential representation of visual motion in the magnocellular system, it is possible that enhancement of the ASNs during saccades may help preserve object motion perception during saccadic sampling of a complex multisensory environment.

The general conclusion discussed above is that the increased BHA just prior to the saccade in the audiovisual movie watching condition reflects amplification of auditory evoked activity coincident with the saccade. An alternative explanation of this BHA increase is that it simply reflects an auditory evoked response to sounds that reliably trigger saccades. Although possible, there are at least two reasons why this explanation is unlikely. First, it relies on the assumption that saccades systematically follow change in auditory activity. To test for this possibility we directly compared the intensity of the auditory stimulus stream before and after fixation onset (+/-0.1, 0.4 and 0.6 sec time intervals) and found no detectable difference. Second, increased BHA was primarily observed on the side contra-lateral to the saccade direction (Fig. S7). This pattern of effects is unlikely to be explained by the evoked response from a salient change to auditory stimulus. In that case one would have expected similar difference on both contra- and ipsi-lateral sides because both ears were stimulated with an auditory stream of comparable qualities (the sound source is in the monitor the subject is watching). Thus the more parsimonious explanation appears to be that a saccade-linked modulatory process increases neural excitability towards the end of fixation and during ensuing saccade.

### Top-down influences of saccade-generating systems

Prior studies suggest that the motor system modulates processing in the auditory system by aligning temporal attention with the timing of predictable events^45–47^. Others have suggested that saccade-generating areas send “corollary discharge/efference copy” signals to sensory areas to prepare them for reafferent input^48^. Our results do not distinguish between these alternatives. It is worth noting, however, that in contrast to covert attention paradigms, where operation of the higher-order mechanism must be inferred from differential performance, tracking the eyes allows us to simply and precisely define an event (the saccade) with clear causal (motor system) antecedents. Thus, while our findings can fit with an attentional explanation (above), they do not require it. Moreover, while there are multiple top-down, bottom-up and lateral circuits conveying visual input into the auditory pathways^35,49^, most neurons in auditory areas do not give frank responses to retinal input; i.e., they do not receive “driving” visual input. These factors lend confidence to our conclusion that saccade effects on processing in classic auditory cortical areas are largely “modulatory” in nature, possibility a primate correlate to effects observed with “head saccades” in the midbrain of the Barn Owl^50^.

During natural active vision, saccade generation provides predictions that are propagated throughout the brain^18^. Our findings suggest that in this process, FEF interactions with the auditory system entail top-down and bottom-up interactions that vary dynamically across the saccade-fixation cycle and are multiplexed in frequency. While this observation is novel for FEF-ASN circuitry, similar multiplexing patterns of connectivity have been reported, including theta-alpha/beta multiplexing in the amygdala-hippocampal circuitry in humans^51^ and prefrontal-hippocampal circuitry in monkeys^52^. Multiplexing in corresponding frequencies has been proposed between several nodes in the visual system with theta and gamma rhythms proposed to serve feed-forward, the beta rhythm proposed to serve feedback influences in monkeys^53^ and humans^54^. Importantly, the current results show that processes tightly linked to saccades transiently organize these multiplexed connectivity patterns modulating both the strength and direction of interaction between the FEF and the auditory system. Our results extend these previous multiplexing connectivity models by showing that network interactions in the alpha and beta ranges may serve interactions in bottom-up as well as top-down directions.

## Author contributions

ML, CES designed the study. ML, SB and MN collected data. ML performed analyses. ML and CES wrote the manuscript. All authors edited the manuscript.

## Methods

### Patients

Continuous eye tracking and iEEG data were recorded from 9 patients (average age 38.6; age range 29; 3 females) implanted with electrodes for surgical treatment of refractory epilepsy. All recordings were performed at the North Shore University Hospital, Manhasset, NY. The study was approved by the institutional review board at the Feinstein Institute for Medical Research and all patients gave written informed consent before implantation of electrodes.

### Task

Participants were presented with a set of 80 images (see Figure 1A for example of an image) each presented for 6 seconds. They were asked to freely explore each image with no constraints or further instructions. After each image participants were asked to rate how much they liked it on a 5 point Likert scale (data not analyzed in the current study). Apart from the free viewing of static images, a subset of patients also performed free viewing of movies with audio (10 min long clip from ‘Despicable Me’ in English; a different 10 min long clip from ‘Despicable Me’ in Hungarian; two repeats of a 4 min long cartoon movie ‘The Present’) and movies with no audio (3 different 5 min long clips of of commercially produced nature documentaries; these data are presented in Fig. 3). All movies were encoded at 30 frames/s. Display size was similar for free viewing of static images movies (∼25-30 dva; see Fig 1A). Similarly to the static images condition, participants were asked to freely explore each image with no constraints or further instructions.

### Electrophysiology data acquisition and preprocessing

Intracranial EEG (iEEG) data were acquired continuously at 1.5 kHz (16-bit precision, range±8mV, DC; Tucker–Davis Technologies, Alachua, FL, USA). Continuous iEEG and eye tracking data were co-registered based on simultaneous time-stamps. Co-registered data were segmented into 6 sec long epochs relative to a stimulus onset and lasting for the duration of stimulus display. Segmented iEEG were down-sampled to 500 Hz for further preprocessing. Subsequently, we removed line noise using band-stop filters (Butterworth 4th order) at 60 Hz, 120 Hz and 180 Hz and created a bipolar montage to maximize spatial specificity of the signal and also to decrease common noise and contributions from distant sources through volume conduction. To this end, we subtracted signals from neighboring contacts on each electrode shaft.

#### Auditory selective channels

We used an auditory localizer to identify auditory selective channels (for a similar approach see for example^55^). These channels were defined as responding with an increased BHA (70-150 Hz) to auditory but not visual stimuli. Participants were passively presented with a set of auditory stimuli (including white noise bursts, syllable trains, music, speech-native and foreign). Subsequently, we compared the amount of BHA elicited by the auditory stimulus relative to the pre-stimulus baseline. To ensure auditory specificity, we only considered channels that responded to auditory stimulus with strong increase from the baseline BHA level (pairwise t-test; t > 10) and did not show any change in response to the onset of the visual image (at the same t-test threshold). This resulted in 220 channels (and 204 after removing channels that showed saccadic spike) across 9 patients. For analysis presented in Fig. S5, we assigned each of these 204 channels into one of 6 anatomical locations. This was done based on individual anatomical landmarks and confirmed by an experienced neurologist. We defined six regions of interest: Heschel’s gyrus (HG; 41 channels, 6 patients), Superior Temporal Gyrus (STG; 69 channels, 9 patients), Frontal Cortex (FC; 28 channels, 6 patients), Parietal cortex (PC; 24 channels, 4 patients), Insular Cortex (IC; 26 channels, 7 patients), Occipital Cortex (OC; 11 channels, 3 patients). Note that the total sum of channels assigned into these six ROIs is 199. The additional 5 auditory selective channels were in the inferior temporal gyrus (2 channels), entorhinal cortex (2 channels) and middle temporal gyrus (1 channel). These five channels were not included in the ROI analyses.

To ensure our results are not driven by activity in channels dominated by polysensory projections from the visual system that terminate in auditory regions^56,57^, we performed a control analysis in which we divided the total pool of channels (N = 204) into pure audio selective channels (N = 155) and mixed audio selective channels (N = 49). Here, we labeled as mixed audio selective all these channels that showed strong auditory response (t_audio_ > 10) in the auditory localizer and weak visual evoked response in the visual localizer (t_visual_ > 1.96 & t_visual_ < 10). Furthermore, we classified as pure audio selective all these channels that showed strong auditory response (t_audio_ > 10) in the auditory localizer and no detectable visual evoked response in the visual localizer (t_visual_ < 1.96). Note that none of the mixed-selective channels showed a classical strong visual evoked potential (see Fig S4-5).

#### Frontal eye fields channels

The frontal eye fields were identified during clinical electrical stimulation mapping using intracortical stimulation (bipolar, symmetric bi-phasic squarewave pulses, 0.2ms/phase, 0.5-4mA, 50Hz for ∼1 sec). Channels were considered to be located in the frontal eye fields if stimulation elicited forced gaze deviation to the contralateral side of stimulation and the anatomical location was in the dorsolateral prefrontal cortex (see^58^ for review). Four patients had pairs of simultaneous recordings in the FEF and auditory system which resulted in the total of 8 FEF channels. Note we performed two analyses of directional network interactions – in one we computed connectivity between FEF and STG channels (127 pairs; Fig. 4); and a control analysis where we computed connectivity between FEF and all ASN channels (232 pairs; Fig. S8).

### Fixation-locked analyses

Spectral phase and power were estimated on data segmented around image presentation (6 sec) with additional 2 seconds before and after image onset and offset to account for edge artifacts of filtering. We used 3 cycles wavelets for frequencies 1-30 Hz in steps of 1 Hz. The complex-valued time series was rectified and squared to extract power. All analysis of neural data during movie watching were performed in the same way as analysis of static images with the same parameters with the exception that an initial segmentation was performed relative to the first and last frame of the movie.

#### Broadband High-frequency Activity (BHA)

was calculated for frequencies 70-150 Hz in 2 Hz steps using a sliding Hanning tapered window (150 ms) with 6 Hz spectral smoothing. The complex-valued signal was rectified and squared to extract power. Frequency dimension was averaged to create a single vector of BHA fluctuations. All analyses were performed on continuous signals before segmenting relative to fixation onset.

Subsequently, we identified time points of fixation onsets and re-segment field potential, time-frequency series and BHA relative to these events. We refer to these fixation-related segments as trials throughout the manuscript. BHA, complex-valued time-frequency series and raw field potentials were re-segmented into 1200 ms long windows with 600 ms before and 600 ms after fixation onset. For further analyses we only considered fixations that were not preceded or followed by another fixation within the +/-200ms window of interest. We defined epochs containing artifacts as those with the gradient of field potential exceeding 5 standard deviations of the trial mean. On average, about 1 % met this criterion and were removed from further analyses.

#### Inter-trial phase coherence (ITC)

was quantified at each time-frequency point. To this end, we used single trial (i.e., fixation locked) time-frequency series complex-valued signal and previously described formula^59^ with the following Matlab implementation:

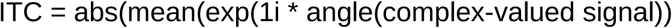

#### Phase Slope Index

To calculate direction of information flux between FEF and auditory selective channels we used Phase Slope Index (PSI^30^). The PSI defines relation between two signals exploiting phase differences across the spectrum. To this end, we used the same preprocessing steps – segmented the data into 6 sec long epochs, band-stop filter to eliminate line noise, create bipolar reference montage to maximize spatial specificity and reduce common sources. Then, we used multitaper frequency transformation with 0.5 Hz frequency resolution on the data padded to 4 sec. Finally, we calculated PSI (with the bandwidth of 2 Hz; i.e., PSI is integrated across 4 frequency points) across all possible pairs of FEF (N = 8) and auditory (N = 53) channels in each of four patients with paired recordings (total number of pairs N = 127). The parameters for spectral smoothing and PSI bandwidth were derived from a simulation study, where we designed two signals one leading the other in the frequency ranges where we expected to observe directional connectivity (i.e., alpha and beta; see simulation study below).

Because PSI is a signed quantity (i.e., positive PSI in the current study means FEF is leading, while negative PSI means STG is leading), the null hypothesis is that PSI scores are drawn from a zero mean distribution^30,60^. To test for significance we used non-parametric Wilcoxon sign rank test with the null hypothesis that the PSI comes from a distribution whose median is zero (N = 127 paired recordings; corrected for multiple comparisons across time and frequency). Note that PSI detects non-zero delays thus being insensitive to volume conduction^30,60^.

#### Statistics

Unless stated differently, we used non-parametric Wilcoxon sign rank test to compare the magnitude spectra phase coherence and power, ERPs and BHA in time windows before and after fixation onset. To test for differences across ROIs we used non-parametric Wilcoxon rank sum test. We used False Discovery Rate (FDR) correction to control for multiple comparisons^61^.

### Eye tracking data acquisition and preprocessing

Eye tracking data were recorded simultaneously with iEEG data. We tracked horizontal and vertical movements of each patient’s left and right eye with a Tobii TX300 eye tracker sampled at 300 Hz. The eye tracker was calibrated by collecting data gaze fixations from five known target points. After the calibration sequence the calibration output was inspected. The calibration output was examined for two features. The calibration points that were either without data or that showed dispersed gaze points scattered around the calibration point were re-calibrated. After calibration was successful, the operator initiated an experiment. Calibration was performed separately for free viewing of static images and for each of the movies. To estimate time-points of fixations and saccades, the position of the eye in x and y coordinates, were transformed into velocities. Velocities exceeding a threshold of 6 times the standard deviation of the velocity distribution for the duration of 12 ms were defined as saccades^62^. The time points of each fixation were further used to analyze perisaccadic field potentials.

