## Supplemental Text and Figures for "Saccadic modulation of neural excitability in auditory areas of the neocortex"

**This PDF file includes:**

Supplementary Text
Figs. S1 to S9

### Supplementary Text

**Could our results reflect extraocular muscle activity rather than genuine neural signals?** Previous studies identified the spike potential in scalp EEG that is generated by extraocular muscle activity[29]. Similar spike potentials have been also found in the intracranial recordings[63,64]. The spectral profile of this spike spares lower frequencies (<30 Hz) but overlaps with BHA range, thus one has to use caution before interpreting the current BHA results. To determine if the BHA signals are contaminated by the extraocular spike, we inspected saccade-locked ERPs (rather than fixation-locked ERPs) in which the spike exceeds neural response by orders of magnitude. The extraocular spike manifests as a bi-phasic transient ERP modulation locked to the onset of the saccade lasting for the duration of the saccade. We identified 16 channels with such a morphology in our data (Fig. S1A-C) primarily in the frontal cortex (N = 8) and anterior portion of the STG (N = 5), anterior insula (N = 2) and one on the border between the insula and Heschel's gyrus (n = 1). Eliminating these channels did not change any of the results (Fig. S1D-G). In particular, the morphology of the BHA power fluctuation, its synchronization with the rhythm of visual exploration (Fig. S1E), as well as overall decreases from the pre- to the post-fixation interval ($z = 2.32$, $p = 0.01$; N = 204) were reproduced after removing these channels. All consecutive analyses are performed on 204 channels which show no detectable sign of extraocular spike potential. While we rule out the possibility of the saccadic spike to explain the current findings, our observations dovetail with previous studies showing that intracranial recordings near the orbits, even with bipolar referencing, are potentially contaminated by extraocular muscle spike potentials[63,64].

**Could our results be explained by weak visual evoked responses?** One possibility is that phase and power perturbations in low frequencies (<30 Hz) noted above reflect fixation-locked visual evoked response, rather than phase perturbation. We defined the ASNs to be auditory-selective but it is possible that some of these channels have weak fixation-locked evoked responses. To test this possibility, we directly compared field potentials in pre- (-100 to 0 ms) and post-fixation intervals (0 to 400 ms in 50 ms bins; Fig S2). To account for possible polarity differences across channels, we first rectified and z-scored fixation locked field potentials. Statistically, ASNs showed no detectable modulation of event related field potentials (all $p > 0.05$ Wilcoxon sign rank test; N = 204; Fig S2; red time-course on the top). However, inspection of responses in each individual electrode, we identified a small fraction of channels in the occipital region that showed an evoked type of response as indexed by post-fixation field potential deviation relative to pre-fixation baseline (Fig. S2; blue line on the bottom). The "multisensory" response profiles in these channels may stem from sparse direct projections to the occipital lobe from the auditory system[65]. To understand whether our results are driven by responses in these sites we performed two control analyses in which we reproduced all our main results with more conservative criteria for classifying electrodes as auditory selective. First, we repeated our analyses after excluding these occipital electrodes, reproducing all our main results without these putative multisensory sites (Fig. S3). To further guard against the possibility that our results depend on activity in similar multisensory channels outside of occipital lobe, we conducted another control analysis in which we identify a subgroup of ASN channels showing what appear to be weak visual evoked potentials (see Methods). This control analysis splits our total pool of channels into pure auditory selective channels (N = 155) and auditory selective channels with strong audio and weak visual evoked response (i.e., mixed channels; N = 49). This control analysis showed that the effects we report do not depend on activity in the mixed auditory selective channels as we reproduced all our main findings in the group of pure auditory selective channels (Fig. S4). It is worth noting that none of these mixed-selective channels showed a classical strong visual evoked potential (see Fig S4D-E).

**Anatomically specific modulations across the ASNs.** To evaluate the anatomical distribution of the effects of saccades on neural activity, we separately studied fixation-locked ITC, low frequency power and BHA in distinct regions of interest (ROIs) across the ASNs. Based on individual anatomical landmarks, we divided all auditory selective channels into six ROIs: Heschel's Gyrus (HG), Superior Temporal Gyrus (STG), Frontal cortex (FC), Parietal cortex (PC), Occipital cortex (OC) and Insular cortex (IC); N = 41, 69, 28, 24, 11, 26 respectively (see Methods for more details). Saccades modulate different aspects of neural activity in each of these areas. Importantly, we found that ITC and power effects noted above had distinct anatomical distributions. While the ITC increase from pre- to post-fixation interval at the rate of saccades was most prominent in the FC, PC and OC, power modulations were most prominent in HG, STG and, PC and IC and OC. Parietal and occipital auditory selective channels intriguingly displayed both of these patterns. ITC without detectable power modulation was primarily observed in FC, while power modulations without detectable ITC was primarily observed in lower level auditory system including HG (Fig S5). These effects were reproduced in a control analyses when we only considered pure auditory selective channels (i.e., after removing channels with weak visual input; Fig. S5A-C).

Despite the variations in ITC and power modulations across ROIs noted above, we also found that BHA magnitude decreased over a period of 75 ms after fixation onset compared to pre-fixation interval in HG ($p = 0.04$; all $z = 2.04$; Wilcoxon sign rank test; $N = 41$). Importantly, at the same time BHA in OC showed an opposite pattern. It increased from pre- to post-fixation interval ($p = 0.04$; $z = 2.04$; Wilcoxon sign rank test; $N = 11$), which is likely explained by an evoked response observed in the OC channels. This suggests that during saccadic sampling of a scene, BHA oscillates with opposite phases in the auditory and visual systems: an increase from pre- to post- fixation in the OC is accompanied by a decrease in other ASNs areas, particularly in the HG. Thus it appears that excitability in the auditory system is suppressed during the time when retinal input is invading occipital cortex. In turn, it is elevated during the saccade, when the OC excitability is reduced. To directly test this possibility that BHA in the auditory and occipital regions is modulated at distinct phases of the saccade-fixation cycle, we directly compared magnitude of BHA in each ROI to that observed in the OC separately in the pre- and post-fixation time windows. In the pre-fixation time interval (-75:0 ms) BHA in all ASN regions was increased compared to OC (all $p < 0.01$; all $z > 2.37$; Wilcoxon rank sum test). In contrast during post-fixation time interval BHA in HG and PC (all $p < 0.006$; all $z > 2.71$; Wilcoxon rank sum test) was decreased compared to the OC. There was no detectable difference between STG, FC, IC and OC (all $p > 0.07$; all $z < 1.79$; Wilcoxon rank sum test). These results show that saccades differently modulate BHA in OC and other ASNs with complementary temporal patterns. While the post-fixation differences may be explained by evoked response in the OC, BHA difference in the pre-fixation time window may reflect differences in the excitability level between OC and other ASN regions. This suggests that before and during saccades, when neural activity in the visual system is suppressed, excitability in the ASNs, as indexed by BHA, increases. In turn, when the visual system shows increased excitability (i.e., after fixation onset), the neural activity in the auditory system (at least in the HG and PC) is reduced. Enhanced auditory excitability before and during saccades may improve sampling of environment when visual system is suppressed (i.e., during saccadic suppression).

**Local activity in the FEF channels.** We identified FEF channels by weak intracortical stimulation (part of the standard clinical mapping procedure) of candidate electrodes localized in the dorsolateral prefrontal cortex. Channels in which stimulation elicited a typical saccadic eye movement to either side were considered as localized in the FEF. We found 8 FEF channels across four patients. Analyzing fixation-locked spectral ITC and power in the FEF channels we observed phase clustering (Fig. 5A) and sustained power above the rate of saccades (7-12 Hz) that extended into beta frequencies (13-30 Hz; Fig. 5B). We noticed that both effects appear modulated by the eye movements, although the comparison was not statistically significant most likely due to our small sample. Next, we evaluated directional network interactions between the FEF and ASNs. Specifically, we focused on the STG channels (54 channels across 4 patients with simultaneous FEF recordings; Fig. 5 C) because this ROI was present in all patients with FEF electrodes and because previous anatomy studies found that neurons in the FEF (particularly the rostral part of the FEF) project primarily to the STG[31]. It is however worth noting that the results are similar when we consider all auditory selective channels rather than STG only (Fig. S8).

**Do our findings reflect the influence of saccades per-se, or that of auditory attention?** The current BHA results may reflect auditory attention, corollary discharge or both. Previous psychophysics studies suggest that as in vision[36], auditory attention is linked to the saccade-fixation cycle in a spatially and temporally specific manner[37-39]. Auditory attention is guided towards a future gaze location even before the movement onset[38]. Importantly, the time course of saccadic influence on auditory discrimination shows that saccade's impact starts prior to movement onset and peaks while the eyes are moving[38] corresponds to the time course of BHA in the current study. These considerations suggest that increased pre-fixation BHA in response to auditory stimulation may have a strong attentional component. To further address this possibility, we examined the spatial specificity of pre-fixation BHA enhancement. We reasoned that if auditory attention is predictively allocated to an upcoming gaze location, we should observe increased BHA on the side *contra-lateral* to the saccade direction in agreement with research showing that contralateral inputs predominate in auditory cortex[66]. In support of this, we observed that auditory stimulation just prior to and that during saccade elicited particularly strong response on the side contra-lateral (both $z = 3.19$; $p < 0.001$; $N = 187$; movies with vs. no audio and movies with audio vs. static images) while it was not significant on the side ipsi-lateral (both $z < 1.37$; $p > 0.16$; $N = 187$) to the saccade direction (Fig. S7A-C). This difference was reproduced in a control analyses in which we excluded all mixed selective channels and limited analyses only to the pure audio selective sites (both $z = 2.79$; $p < 0.005$; $N = 145$; movies with vs. no audio and movies with audio vs. static images; Fig S7D-F). Further analyses showed that the effect was primarily visible in the STG ($z = 2.78$ ; $p = 0.005$; $N = 49$) but not in other ROIs (all $z < 1.53$; all $p > 0.12$). These results support the model in which auditory attention is linked to the saccade-fixation cycle in spatio-temporal manner. Auditory attention amplifies neural responses to auditory stimulation late in fixation period and during the ensuing saccade primarily on the side contralateral to the upcoming saccade direction.

**What is the optimal frequency integration window for the PSI: a simulation study.**
PSI requires calculating a slope across multiple frequencies. Several possible frequency windows may be considered. We run simulations to identify an optimal bandwidth across which the slope should be integrated given frequencies where we hypothesized our effects to be strongest. To this end, we generated two signals (1 sec long sampled at 500 Hz) in a way that one is a copy of the other shifted in time by 50 milliseconds. This temporal lag is within the physiological range of excitability modulations from the FEF to sensory system[67]. Because the highest frequency range in which we expected network interactions between FEF-ASNs is in the beta range (e.g., 13-17 Hz[46]) we created oscillations that were centered at this frequency range. We added Gaussian noise to each of these signals and repeated this procedure 100 times to approximate 100 trials. Next, we calculated PSI using the established method (see above). To identify optimal bandwidth for the PSI we systematically explored the bandwidth frequency window from 1 to 10 Hz. As expected we noted a trade-off between the window size and frequency precision. We also observed there was no difference in the estimated PSI magnitude for windows over 2-3 Hz (see Fig. 9A-B). However, their frequency resolution and accuracy of frequency estimation was gradually decreasing. To optimize the trade-off between window size and frequency resolution we selected window size of 2 Hz (i.e., 4 frequency points at 0.5 Hz frequency resolution). This selection was also confirmed in another simulation in which we created

similar signals with the same lag. However, now we introduced a multiplexed connectivity pattern between these two signals[46] - signal 1 was leading in theta (3-7 Hz) and lagging in beta frequency (13-17 Hz; Fig. S9B). Altogether, these simulations convinced us that a frequency window of 2 Hz (i.e., 4 frequency points) may be an optimal for the purpose of testing the current hypothesis. It is worth noting that another study found the same bandwidth to be optimal for studying connectivity in a similar frequency range[68]. Finally, to further ensure that our main results are robust across bandwidth parameters we re-calculate the PSI between FEF and STG with the bandwidth of 6 Hz (i.e., 12 frequency points). This analysis reproduced all our main effects and conclusions (Fig. S9C).

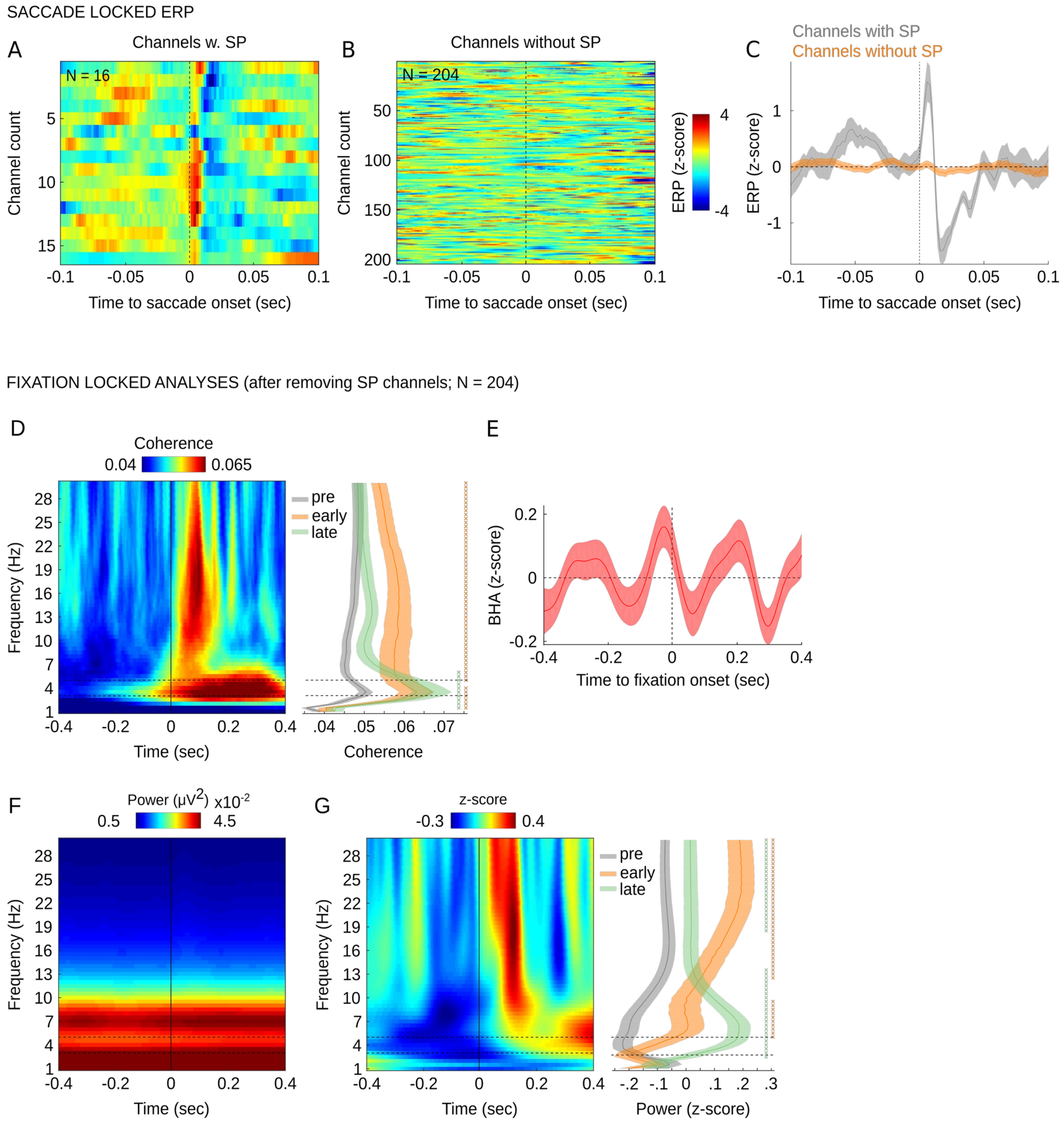


***Figure S1. Saccade-locked activity to identify saccadic spike potential.*** *We identified 16 channels with a bi-phasic transient ERP modulation locked to the onset of the saccade lasting for the duration of the saccade (aka saccadic spike potential; A) and 204 channels with no detectable saccadic spike potential (B).* ***(A, B)*** *Color maps show ERPs locked to the saccade onset for channels with (A) and without (B) saccadic spike potential. X-axis show time relative to the saccade onset, y-axes show*

*individual channels. Color indicates ERP magnitude.* ***(C)*** *ERPs with (gray) and without (orange) saccadic spike potential averaged across channels. In control analyzes (D-G) we reproduce our main results presented in Fig 2 of the main manuscript for 204 channels without the saccadic spike potential.* ***(D)*** *Color map shows grand average fixation-locked ITC (N = 204 ASN channels without the saccadic spike potential, 9 patients; time on x-axis, frequency on y-axis). Vertical line indicates fixation onset. Horizontal dashed lines bracket the frequency range of saccadic eye movements (i.e., 3 – 5 Hz). Right panel shows frequency distributions of phase coherence averaged within three time windows relative to fixation-onset: pre-fixation (gray; -400:0 ms), early-post (0-100 ms; orange) and late-post (100:400 ms; green). Shading reflects standard error of the mean (SEM). Horizontal dashed lines bracket the frequency range of saccadic eye movements.* ***(E)*** *Line plot shows z-scored BHA (70-150 Hz) locked to fixation onset aggregated across all 204 auditory selective channels without the saccadic spike potential.* ***(F, G)*** *Color map shows fixation locked power without normalization (F) and normalized (G) by z-scoring each frequency separately to remove the 1/f component. In both panels time is on x-axis and frequency on y-axis. As in panel A, horizontal dashed lines bracket the frequency range of saccadic eye movements and the vertical solid line indicates the time of fixation-onset. The right panel shows frequency range of power fluctuations within three time windows relative to fixation-onset (as in panel D). All results are controlled for multiple comparisons with Benjamini & Hochberg/Yekutieli false discovery rate procedure. "X" symbols in right panels A, D indicate p-values < 0.05 separately for the comparison between pre- vs. early-post fixation and pre- vs. late-post fixation time window (orange and green, respectively).*

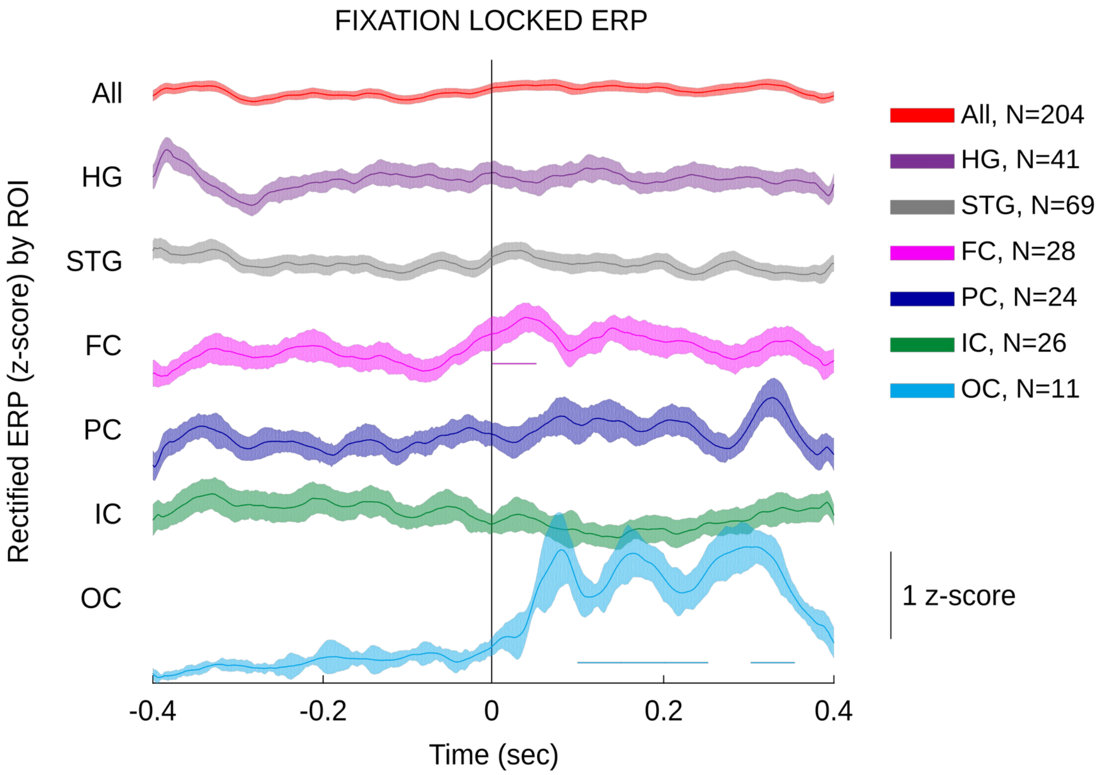


***Figure S2. Fixation-locked evoked field potentials across ROIs.*** *Fixation-locked ERPs rectified and z-scored are plotted separately for each Region of Interest (ROI): Heschel's gyrus (HG), Superior Temporal Gyrus (STG), Frontal cortex (FC), Parietal cortex (PC), Insular cortex (IC) and Occipital cortex (OC). Solid line reflects mean ERP for each ROI. Shading reflects standard error of the mean. Line symbols below FC and OC indicate p-values < 0.05. This is a result from a Wilcoxon sign rank test comparing post-fixation intervals (i.e., 8 consecutive intervals from 0 to 400 ms post-fixation each lasting 50 ms) relative to pre-fixation baseline (-100:0 ms). Note that the earliest interval in FC and several later periods in OC ERPs show saccade-related modulation but no other ROI shows detectable modulation in the ERPs.*

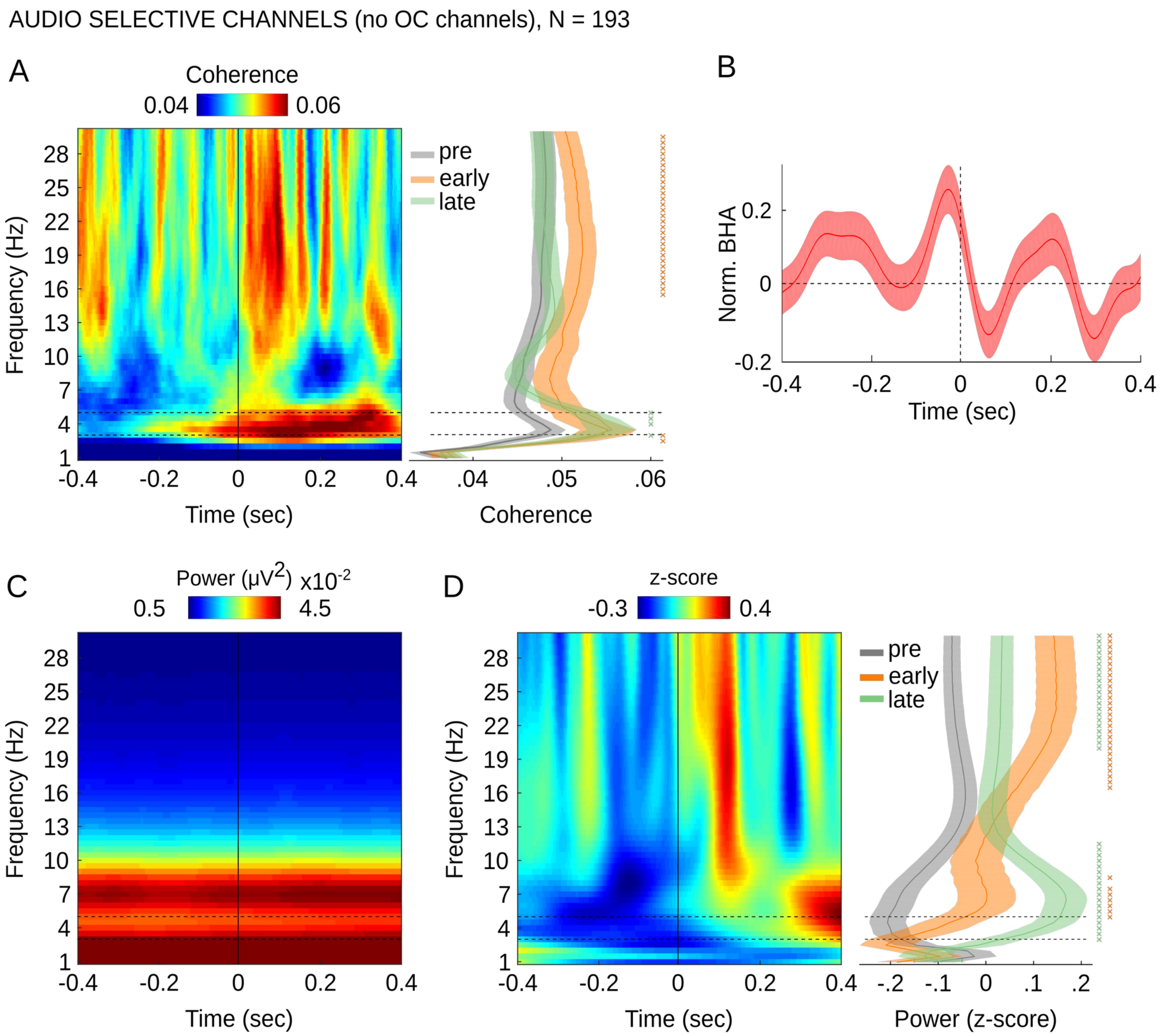


***Figure S3. Fixation-locked neural activity in the auditory selective channels after removing occipital channels**. We observed that channels localized in the occipital lobe but no other ROI showed an evoked response (see Fig S2). In a control analysis, we removed all these channels and repeated all analyses presented in Figure 2. **(A)** Grand average fixation-locked phase concentration (N = 209 ASN channels, 9 patients; no occipital channels). Color map shows inter-fixation phase coherence (time on x-axis, frequency on y-axis). Vertical line indicates fixation onset. Horizontal dashed lines bracket the frequency range of saccadic eye movements (i.e., 3 – 5 Hz). The right panel shows frequency distributions of phase coherence averaged within four time windows relative to fixation-onset: pre-fixation (gray; -400-0 ms) post-fixation (0-400 ms; orange), early-post (0-100 ms; violet) and late-post (100-400 ms; green). Shading reflects standard error of the mean (SEM).*

*Horizontal dashed lines bracket the frequency range of saccadic eye movements. Marks on the right side indicate significant phase coherence change in each of the post-fixation windows relative to the pre-fixation interval.* ***(B)*** *Upper panel line plot shows z-scored BHA (70-150 Hz) locked to fixation onset (left panel) aggregated across all 209 auditory selective channels (no occipital channels). Lower panel shows the result of spectral analyses using the Fast Fourier Transform of the BHA. The peak in spectrum at the rate of eye movements shows that the BHA in ASNs oscillates at the rate of eye movements. X-axis show frequency and y-axis show the magnitude of BHA modulation.* ***(C)*** *Color map shows fixation locked power with no normalization (left panel) and normalized by z-scoring each frequency separately to remove 1/f (middle panel). In both panels time is on x-axis and frequency on y-axis. As in panel A, horizontal dashed lines bracket the frequency range of saccadic eye movements and the vertical solid line indicates the time of fixation-onset. The right panel shows frequency distributions of power within four time windows relative to fixation-onset (same as in panel A and B). Marks on the right side in panel A and C indicate significant phase coherence (A) and power (C) change in each of the post-fixation windows relative to pre-fixation time interval. All results are controlled for multiple comparisons with Benjamini & Hochberg/Yekutieli false discovery rate procedure. Star symbols in panels A-C indicate p-values: * reflects $p < 0.05$, ** reflects $p < 0.01$, *** reflects $p < 0.005$.*

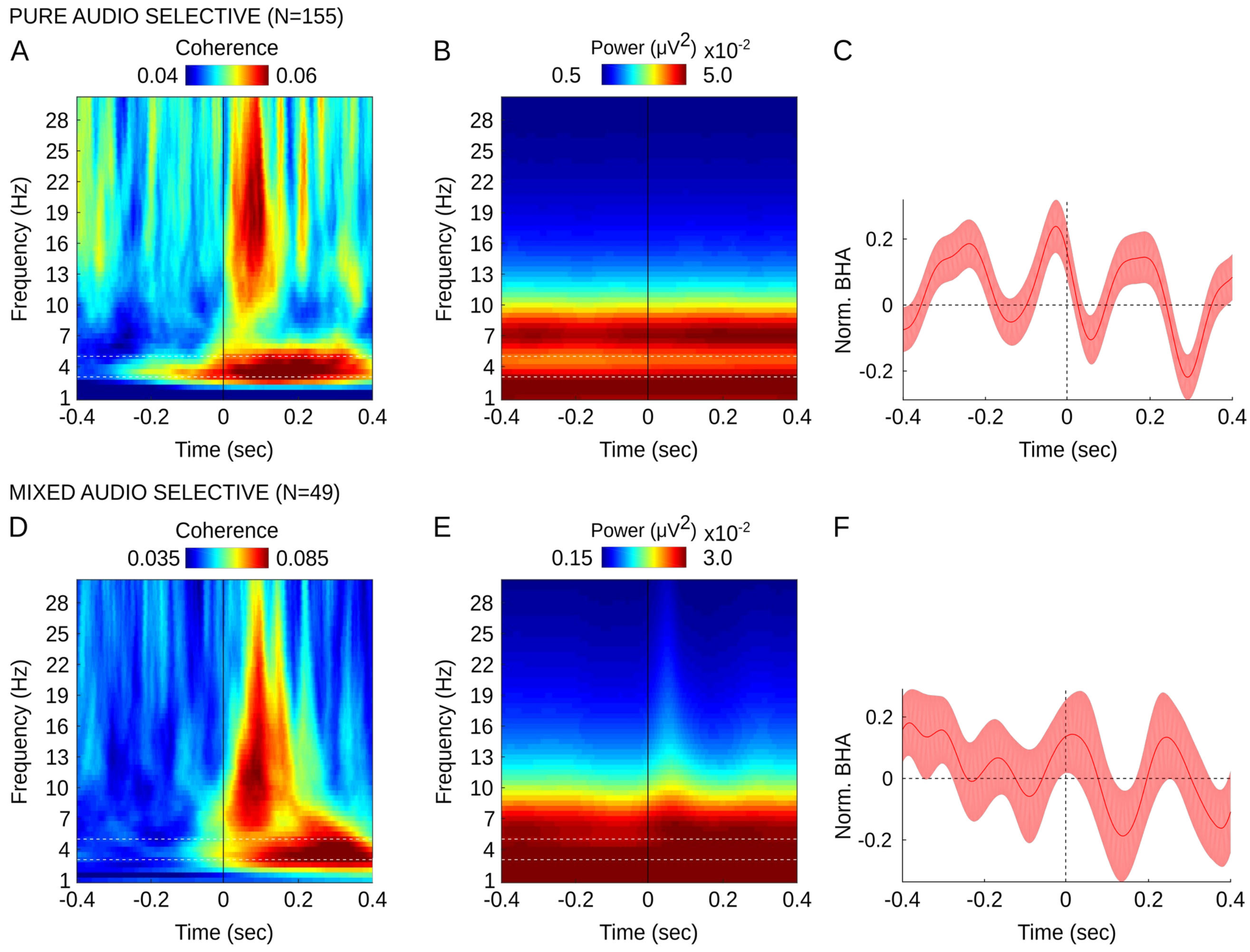


***Figure S4. Fixation-locked neural activity in pure auditory selective channels vs. mixed auditory selective channels.*** *In a control analysis we split the total pool of 204 audio selective channels into 155 pure audio selective channels (i.e., channels that showed strong auditory response with t > 10 – and no sign of visual evoked response at the lowest threshold with t < 1.96) and 49 mixed audio selective channels (i.e., channels that showed strong auditory response with t > 10 and weak visual evoked response with t < 10). The purpose of this analyses was to assure that the results are consistent across subpopulations of ASN channels. Furthermore, we wanted to assure that the results are also consistent when we exclude mixed audio selective channels that might show weak visual evoked potentials. Results are consistent with these reported in Fig 2 of the main manuscript. Color maps show ITC (A,D) and power (B,E) in pure auditory selective channels (A,B) and mixed auditory selective channels (i.e., containing weak visual input bias; D,E). X-axes show time relative to fixation onset, Y-axes show frequency. The vertical line indicates fixation onset. Horizontal dashed lines bracket the frequency range of saccadic eye movements (i.e., 3 – 5 Hz). Line plots* ***(C, F)*** *show shows z-scored BHA (70-150Hz) locked to fixation onset in in pure auditory selective channels (C) and*

*mixed auditory selective channels (F). X-axis show time relative to fixation onset, Y-axes show BHA magnitude. Vertical line indicates fixation onset. Note that the peak timing of the fixation-BHA signal for the auditory channels with putative visual input has shifted to a pattern more like that of a visual area (post-fixation increase in excitability).*

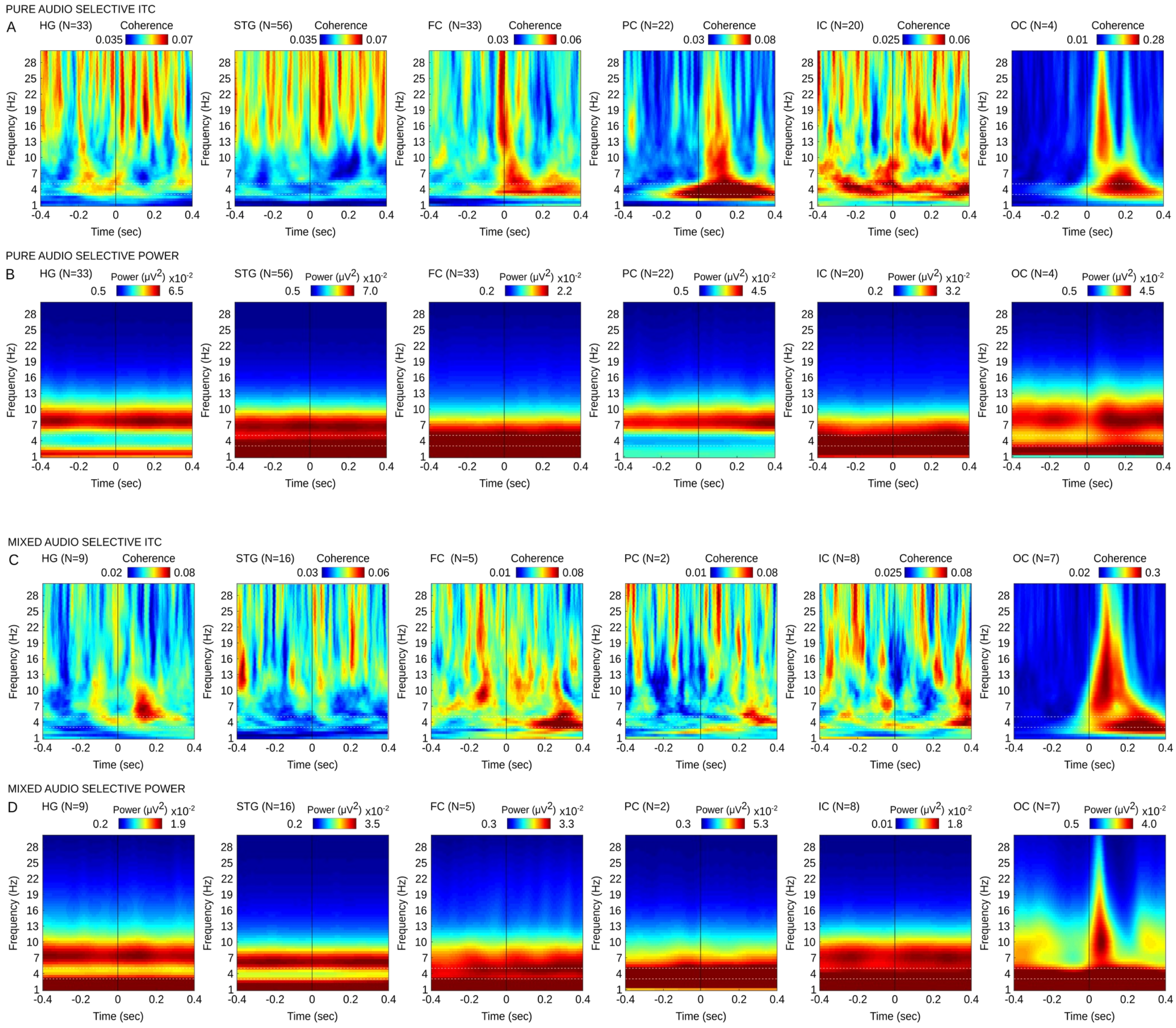


***Figure S5. Fixation-locked ITC and power in pure auditory selective channels and mixed auditory selective channels.*** *In a control analysis we tested whether the effects of ITC and power noted in separate ROIs are preserved n both pure audio selective channels (i.e., channels that showed strong auditory response – t > 10 – and no sign of visual evoked response at the lowest threshold – t = 1.96) and mixed audio selective channels (i.e., strong auditory response with t > 10 along with weak visual evoked response having t < 10 & t > 1.96). The purpose of this analyses was to assure that the results are consistent across subpopulations of ASN channels. Furthermore, we wanted to assure that the results are also consistent when we exclude mixed audio selective channels that might show weak visual evoked responses. Briefly, results are consistent with these reported in Fig 2 of the main paper. Rows A and C show ITC in pure audio selective channels* ***(A)*** *and mixed audio selective channels* ***(C).*** *Rows B and D show power in pure audio selective channel* ***(B)*** *and*

*mixed audio selective channels* ***(D).*** *Each color map shows data from a single ROI:  Heschel's gyrus (HG), Superior Temporal Gyrus (STG), Frontal cortex (FC), Parietal cortex (PC), Insular cortex (IC) and Occipital cortex (OC). Note that the total number of ASN channels 220 includes 215 channels in six ROIs and 5 other channels that did not belong to any of these ROIs (see Methods).*

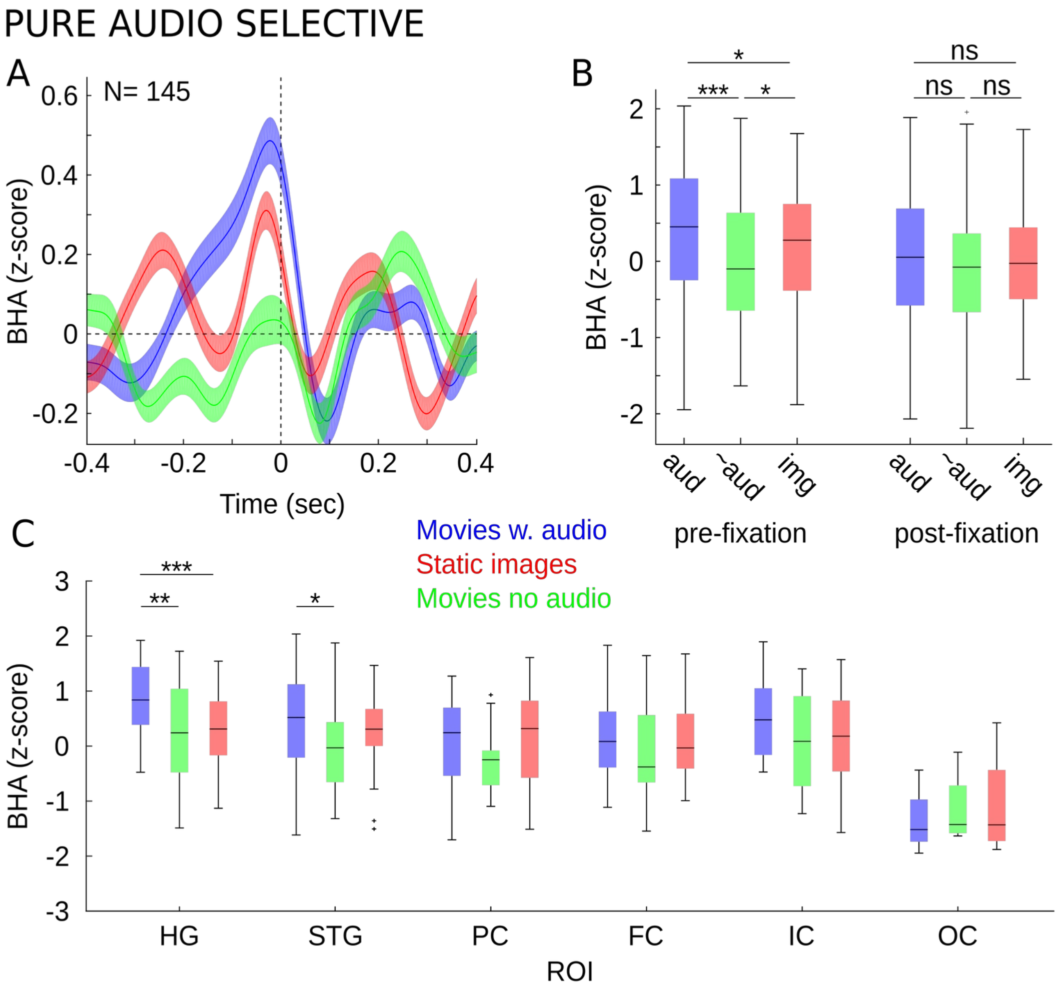


***Figure S6. Fixation-locked BHA in three viewing conditions after removing mixed selective channels.*** *In a control analysis we studied fixation-locked BHA in 145 pure audio selective channels (i.e., channels that showed strong auditory response – t > 10 – and no sign of visual evoked response at the lowest threshold – t = 1.96) in three viewing conditions: free viewing of movies with an audio stream (blue), without the audio stream (green) and free viewing of static images (red). Results are consistent with these reported in Fig 4 main manuscript. Panel* ***(A)*** *shows the time course of fixation-locked BHA in these three conditions. X-axis - time relative to fixation onset; Y-axis - magnitude of normalized BHA. Normalization was performed by z-scoring BHA in each condition across time. Box plots in panel* ***(B)*** *show magnitude of BHA in three viewing conditions separately for the time interval before fixation onset (left panel; -100 ms to 0 ms) and after-fixation onset (right panel; 0 ms to 100 ms). Y-axis - magnitude of BHA. Box plots in* ***(C)*** *show pre-fixation BHA in the three viewing conditions separately for each ROI: Heschel's gyrus (HG), Superior Temporal Gyrus (STG), Frontal cortex (FC), Parietal cortex (PC), Insular cortex (IC) and Occipital cortex (OC). Box plots in (B and C) indicate 25$^{th}$, median and 75$^{th}$ percentile, whiskers extend to extreme values not considered outliers while outliers*

*are marked with crosses. Star symbols indicate p-values: * reflects $p < 0.05$; ** reflects $p < 0.01$; *** reflects $p < 0.005$; ns reflects $p > 0.05$. In this control analysis fixation-locked BHA was increased prior to fixation onset in movies with audio as compared to both movies with no audio ($z = 4.22$, $p < 0.001$; $N = 145$) and free viewing of static images ($z = 2.20$; $p = 0.02$; $N = 145$) while we found no difference in the post-fixation time window (all $z < 1.6$; $p > 0.10$; $N = 145$). This pre-fixation enhancement was detectable in HG and STG (both $z > 2.43$; both $p < 0.01$; $N = 145$; movies with audio vs. movies with no audio; Fig. S6C), but not in other ROIs (all $z < 1.57$; all $p > 0.11$).*

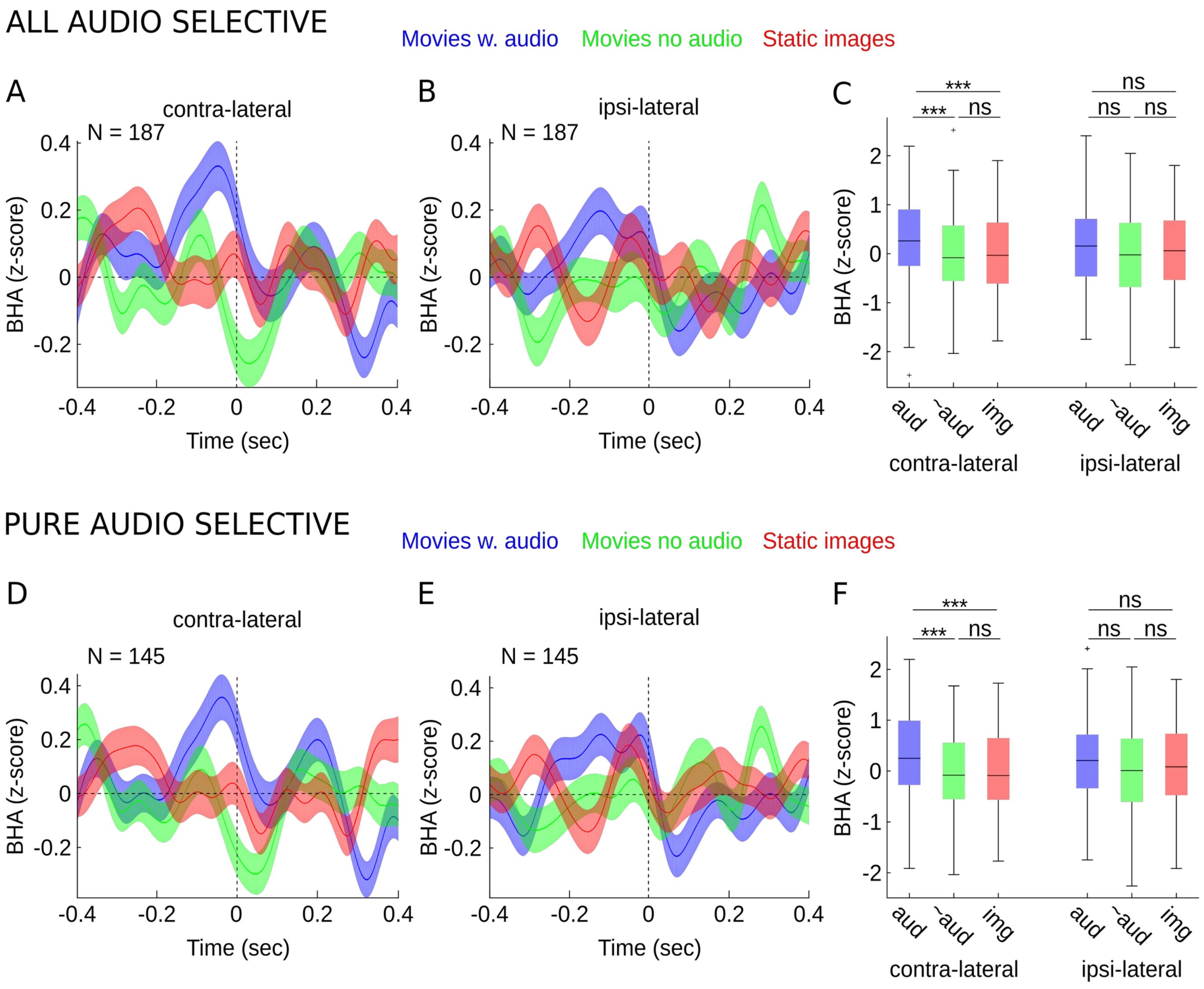


***Figure S7. Fixation-locked BHA in three viewing conditions separately for contra- and ipsi-lateral saccade direction in all (A-C) and pure audio selective channels (D-F). (A, B, D, E)*** *The time course of fixation-locked z-scored BHA time course in three viewing conditions: free viewing of movies with audio stream (blue), movies with no audio (green) and static images (red) for all audio selective channels (A, B) and pure audio selective channels (D, E). For each electrode the data are locked to fixation onset that follow a saccade towards contra-lateral direction (A, D) and ipsi-lateral direction (B, E). X-axis show time relative to the fixation onset. Y-axes show magnitude of normalized BHA. Normalization was performed by z-scoring BHA in each condition across time. Box plots in panel* ***(C, F)*** *show magnitude of BHA in three viewing conditions for time interval before fixation onset (-100 ms to 0 ms) separately for saccades towards contra- (left panel) and ipsi-lateral (right panel) direction. Box plots indicate 25$^{th}$, median and 75$^{th}$ percentile, whiskers extend to extreme values not considered*

outliers while outliers are marked with crosses. Star symbols indicate p-values: * reflects $p < 0.05$; ** reflects $p < 0.01$; *** reflects $p < 0.005$; ns reflects $p > 0.05$.

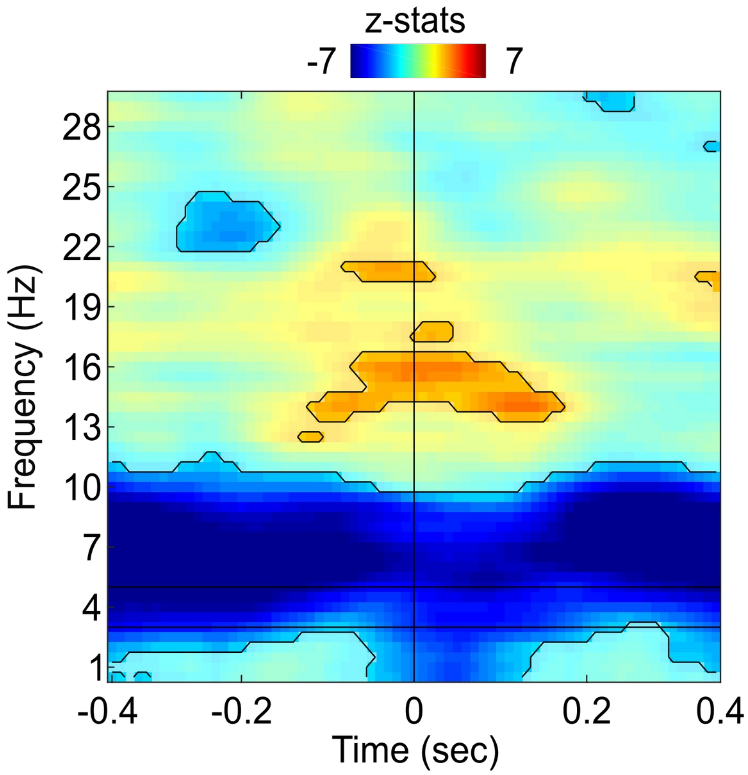


***Figure S8. Fixation-locked directional network connectivity between FEF and all ASN channels.*** *Color map shows fixation-locked z-statistics from sign-rank test comparing PSI against a null hypothesis (i.e., PSI not different from zero). Negative z-statistics values indicate time-frequency points when ASNs lead FEF (i.e., bottom-up directional flow), positive values indicate points when FEF leads ASNs (i.e., top-down directional flow). Vertical line marks fixation onset. Horizontal dashed lines show lower and higher cutoff frequencies of saccadic eye movements. Contours depict significant time-frequency points ($p < 0.05$ controlled for multiple comparisons).*

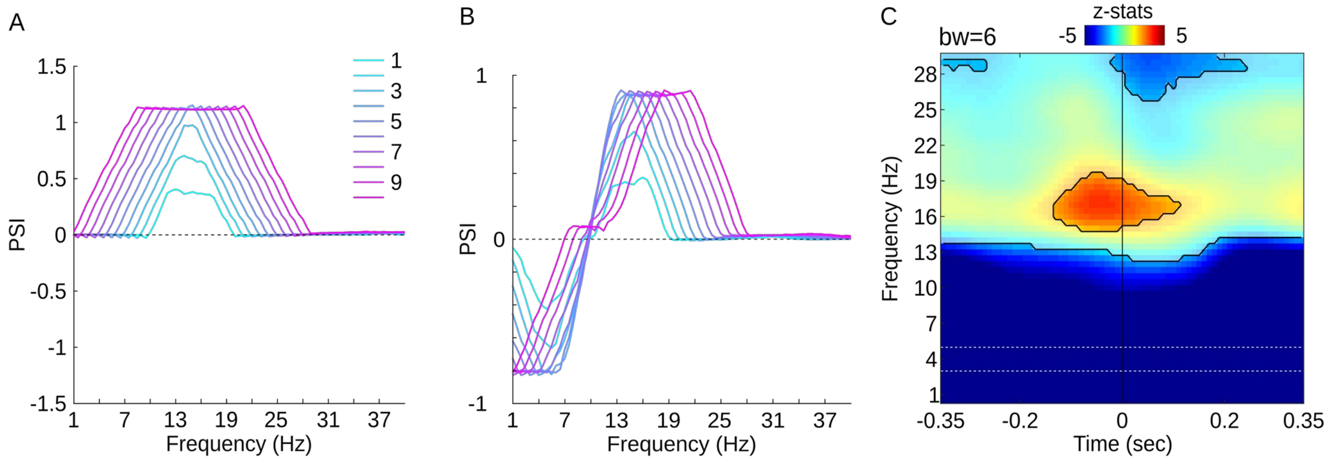


***Figure S9. Simulation study to define optimal bandwidth for the PSI and a control analysis with PSI between FEF and STG calculated with increased bandwidth of 6 Hz.***

***(A)*** *Results from a simulation show PSI between two signals where signal-1 leads signal-2 in beta frequency (13-17 Hz). Color lines show PSI calculated for linearly increasing bandwidth (1-10 Hz).* ***(B)*** *Same as A but simulated signals show a multiplexed connectivity profile. Signal-1 leads signal-2 in beta frequency (13-17 Hz; positive PSI) and signal signal-1 lags signal-2 in theta frequency (3-7 Hz; negative PSI).* ***(C)*** *Here, we used an increased bandwidth of 6 Hz to guard against a possibility that the PSI results observed in the current study depend on selected bandwidth. Color map shows fixation-locked z-statistics from a sign-rank test comparing PSI (bandwidth of 6 Hz) against a null hypothesis (i.e., PSI not different from zero). Negative z-statistics values indicate time-frequency points when STG leads FEF (i.e., bottom-up directional flow), positive values indicate points when FEF leads STG (i.e., top-down directional flow). Vertical line marks fixation onset. Horizontal dashed lines show lower and higher cutoff frequencies of saccadic eye movements. Contours depict significant time-frequency points ($p < 0.05$ controlled for multiple comparisons). Briefly, all effects are comparable to that observed in the main manuscript.*
